# Insights into the role of phosphorylation on microtubule crosslinking by PRC1

**DOI:** 10.1101/2024.10.04.616702

**Authors:** Ellinor Tai, Austin Henglein, Angus Alfieri, Gauri Saxena, Scott Forth

## Abstract

The mitotic spindle is composed of distinct networks of microtubules, including interpolar bundles that can bridge sister kinetochore fibers and bundles that organize the spindle midzone in anaphase. The crosslinking protein PRC1 can mediate such interactions between antiparallel microtubules. PRC1 is a substrate of mitotic kinases including CDK/cyclin-B, suggesting that it can be phosphorylated in metaphase and dephosphorylated in anaphase. How these biochemical changes to specific residues regulate its function and ability to organize bundles is not known. Here, we perform biophysical analyses on microtubule networks crosslinked by two PRC1 constructs, one a wild-type reflecting a dephosphorylated state, and one phosphomimetic construct with two threonine to glutamic acid substitutions near PRC1’s microtubule binding domain. We find that the wild-type construct builds longer and larger bundles that form more rapidly and are much less resistant to mechanical disruption than the phosphomimetic PRC1. Interestingly, microtubule pairs organized by both constructs behave similarly within the same assays. Our results suggest that phosphorylation of PRC1 in metaphase would tune the protein to stabilize smaller and more flexible bundles, while removal of these PTMs in anaphase would favor the assembly of larger more mechanically robust bundles to resist chromosome and pole separation forces at the spindle midzone.

## Introduction

Cell division in eukaryotes requires the proper assembly of the mitotic spindle^1,2^. The spatiotemporal organization of this complex microtubule network occurs via an interplay of diverse motor and non-motor protein factors needed to properly align and segregate chromosomes, orient the spindle, and position the cell division plane^2,3^. Within the spindle, these networks include kinetochore fibers which make attachments to chromosomes, astral microtubules that interact with the cell cortex, and interpolar microtubules that span the length of the spindle, form the bulk of the spindle microtubule mass, and provide structural robustness^4,5^. Near the center of the spindle, interpolar microtubules interdigitate with overlapping plus-ends to form antiparallel arrays, providing a platform for motor proteins such as kinesin-5 to slide microtubules outward towards the poles^6,7^. Recent evidence suggests that in metaphase a sub-population of interpolar microtubules form bundles which connect to sister kinetochore fibers at their minus-ends^8–12^. Termed “bridging fibers” these bundles are proposed to provide a mechanical coupling of k-fibers while spanning the center of the spindle and helping to regulate chromosome centering motions^9–12^. Bridging fibers then persist into anaphase, evolving to form the larger bundles of the spindle midzone whose microtubules are far less dynamic and feature reduced overlap lengths^12–15^.

The organization of this specialized subset of interpolar microtubules has been shown to depend on the microtubule crosslinking activity of protein regulator of cytokinesis-1 (PRC1)^16–18^. Like other homologs of the MAP65 family of proteins, which include Ase1 in yeast and MAP65 in plants^19–21^, PRC1 preferentially crosslinks microtubules in an antiparallel orientation and is particularly enriched within the spindle midzone in anaphase^22–25^. Depletion or optogenetic deactivation of PRC1 results in chromosome alignment defects in metaphase, reduced coupling between sister kinetochore motions, and an increase in lagging chromosomes in anaphase^26^. Microneedle manipulation experiments in PRC1-depleted metaphase cells demonstrated that bridging fibers provide a mechanical coupling which transmits pulling and bending forces between sister kinetochore fibers^27^. Extensive *in vitro* experiments have also provided evidence that PRC1/Ase1 mechanically couple microtubule pairs^28,29^. We have previously shown via *in vitro* reconstitution experiments that PRC1 ensembles can act as viscous dashpots, providing velocity-dependent resistance to microtubule sliding^30^. Additionally, we reported that when subjected to kinesin-driven sliding forces, a distinct braking mode emerges when PRC1 molecules are densely clustered near microtubule tips^31^. The yeast homolog, Ase1, has been shown to act as an adaptive brake against sliding by kinesin-14 motor proteins^32,33^, as well as providing resistance against filament separation via a unique entropic expansion mechanism^28^.

The function and localization of mitotic proteins is precisely regulated by post-translational modifications, particularly the kinase activity of CDK/cyclin-B^16^. In metaphase, when CDK/cyclin-B levels and activity are high, numerous proteins are phosphorylated. Upon entry into anaphase, cyclin-B levels rapidly decrease and phosphorylation marks are removed from many substrates, likely altering their function and localization^34–36^. PRC1 has been shown to be a substrate of multiple kinases, including polo-like kinase 1^37^ and CDK/cyclin-B^34^, which regulate its recruitment to microtubules as well as multiple binding partners required for positioning the cell division plane. In particular, two threonine residues within PRC1 at positions 470 and 481 were confirmed to be targeted for phosphorylation by CDK/cyclin-B^38^. Phosphorylation-null T470/481A mutants were shown to significantly enhance microtubule bundling in pro-metaphase, suggesting that phosphorylation at these residues tunes bundling to be less robust throughout metaphase^35^. These residues are of special interest because they are located within an unstructured C-terminal tail domain just adjacent to PRC1’s microtubule binding spectrin domain, suggesting that modifying the charges near the negatively charged microtubule lattice can play an important role in microtubule crosslinking and bundling, especially within metaphase bridging fibers when PRC1 is likely phosphorylated at these residues^24^. As PRC1 undergoes dephosphorylation upon entry into anaphase, its microtubule bundling properties likely change. However, it is currently unclear how these PTMs alter the kinetics of bundle formation and the mechanics of bundle stability under load.

To address these outstanding questions, we used site-directed mutagenesis to generate a phosphomimetic construct mimicking PRC1’s state while phosphorylated by CDK/cyclin-B (T470/481E) to directly compare fundamental biophysical properties of the protein against the wild-type dephosphorylated construct we have previously reported on^30,31^. We employed a series of reconstitution and motor-driven gliding assays along with total internal reflection fluorescence (TIRF) microscopy to characterize the microtubule bundling properties of these two PRC1 constructs both at and far from mechanical equilibrium. We found that phosphomimetic PRC1 assembled shorter bundles containing few microtubules than the wild-type dephosphorylated construct, and the rate of assembling these structures was significantly slower. Interestingly, the assembly and stability of pairs against outward sliding forces were similar for both constructs, but the disassembly of higher order bundles containing >4 microtubules occurred much more rapidly and completely for the phosphomimetic construct. Structural predictions of the effect of the two amino acid substitutions suggest that subtle conformational changes near the microtubule surface might propagate along the length of the PRC1 dimer^39^, leading to a dramatic change in the ability to assemble multi-filament micron-scale networks. Together, our results suggest that PRC1 likely generates two distinct types of bundles in metaphase and anaphase that are optimally tuned for the unique functions of interpolar microtubules in each of these phases of mitosis.

## Results

### T470/481E-PRC1 assembles shorter bundles that contain fewer microtubules than WT-PRC1

We sought to understand how the mechanics of microtubule crosslinking by PRC1 might differ between metaphase and anaphase. PRC1 is well-known to crosslink microtubules in an antiparallel configuration (Fig. 1A) and localize to the spindle interpolar microtubule network. In metaphase, when CDK/cyclin-B levels are high, PRC1 has been shown to be phosphorylated at two threonine residues located at sites 470 and 481 within a largely unstructured tail region in the C-terminus, just adjacent to the microtubule-binding spectrin domain (Fig. 1B). Upon entry into anaphase, these phosphorylation marks are removed. We therefore wished to generate and purify protein constructs reflecting these two different states (Fig. 1B). We have previously reported on the expression of a wild-type construct^30,31^ that reflects PRC1 in the anaphase state. We performed site-directed mutagenesis to mutate the two threonine residues at sites 470 and 481 to glutamic acids to generate a phosphomimetic construct that mimics the metaphase state of PRC1. We henceforth refer to these two constructs as wild-type PRC1 (WT-PRC1) and phosphomimetic PRC1 (T470/481E-PRC1).

**Figure 1.**
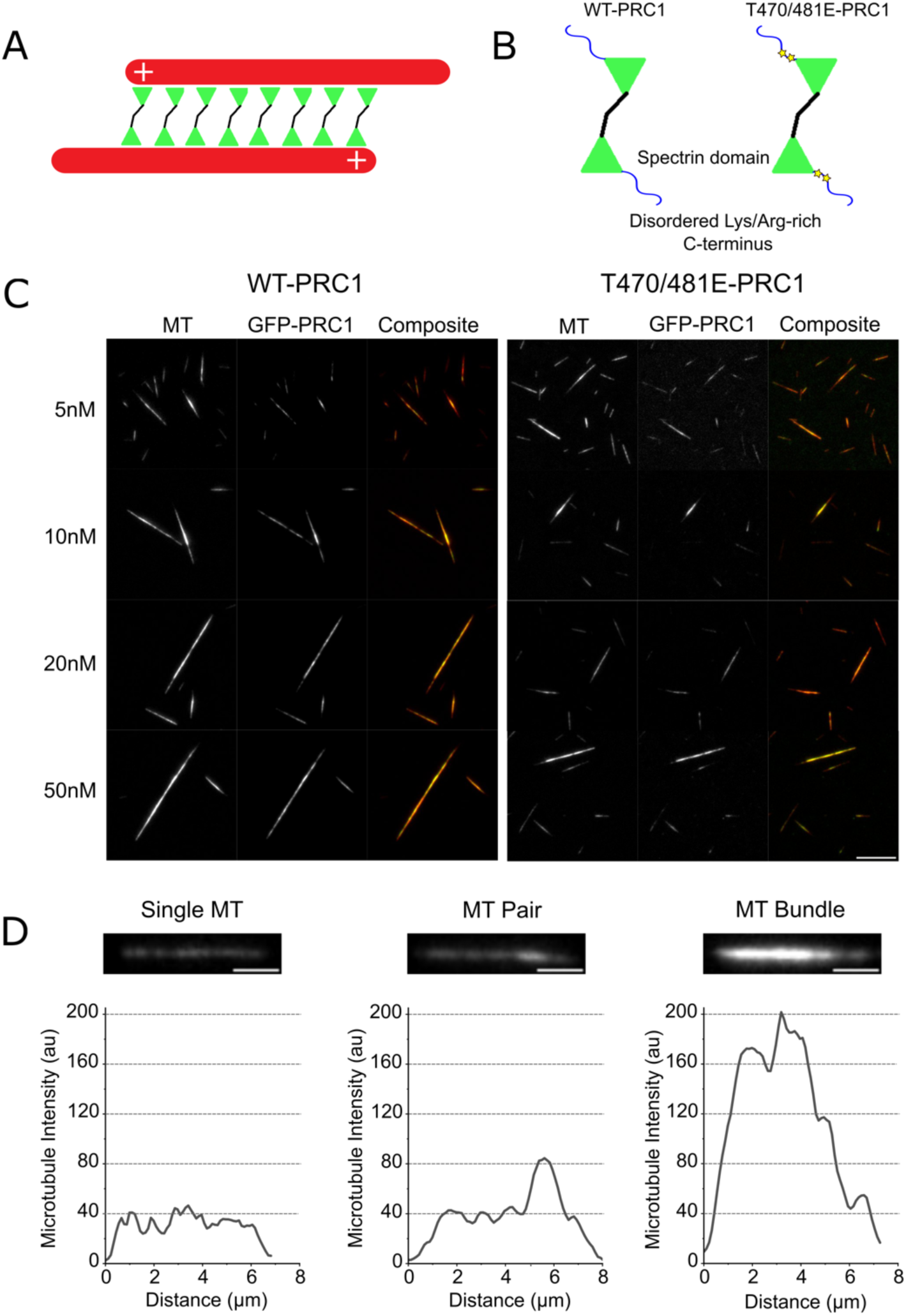
PRC1 crosslinks microtubules into pairs and higher order bundles. (A) A schematic depicting PRC1 molecules crosslinking two antiparallel microtubules. (B) Domain schematic depicting location of residues T470 and T481, which were replaced with phosphomimetic E residues to generate GFP-T470/481E-PRC1 for these studies. (C) Fluorescence images of microtubules (red) and GFP-PRC1 (green) forming bundled structures after incubation for 10 minutes at different solution concentrations of PRC1 (5, 10, 20, 50nM respectively). Scale bar = 10 µm. (D) Examples of microtubule single, pair, and bundle with corresponding intensity linescan. Scale bar = 2 µm.

We next wanted to determine the extent to which PRC1 could bundle microtubules. We incubated GMPCPP-polymerized and taxol-stabilized microtubules labeled at a 1:100 ratio with rhodamine tubulin with purified GFP-PRC1 molecules at different concentrations. After allowing microtubules and PRC1 to equilibrate, we deposited the resulting structures on the surface of a coverslip and imaged both microtubules and GFP-PRC1 using 2-color TIRF microscopy. Representative images across a range of PRC1 concentrations (5, 10, 20, 50 nM) for both WT-PRC1 and T470/481E-PRC1 were collected (Fig. 1C) and a variety of differing size, microtubule number, and PRC1 content were observed. To quantify the number of microtubules contained within each individual object, we performed linescan analysis of the rhodamine fluorescence signal along the length of the structure, and were readily able to distinguish between single microtubules, microtubule pairs, and bundles containing three or more microtubules (Fig. 1D).

With these calibrations, we were next able to characterize key properties of the bundles to identify differences in equilibrium crosslinking between the two PRC1 constructs. We first measured the average number of microtubules found per structure at each of the measured PRC1 concentrations. For WT-PRC1, the average number of microtubules within a structure was ∼3 at 5nM PRC1, which increased up to nearly 6 microtubules per bundle at 50nM PRC1 (Fig. 2A). In contrast, for the T470/481E construct, bundles of approximately the same size (∼3) formed at 5nM, but did not significantly increase beyond this size even up to 50nM of PRC1 added (Fig. 2A). We next examined the lengths of individual bundled structures, and found a similar relationship. When bundling with WT-PRC1, bundles were longer than when using T470/481E-PRC1 at the same concentration, and increasing PRC1 concentration yielded longer bundles for the WT-PRC1 but exhibited little length increase for T470/481E-PRC1. Together, these results suggest that phosphomimetic PRC1 is less efficient at forming multi-microtubule bundles relative to wild-type PRC1.

**Figure 2.**
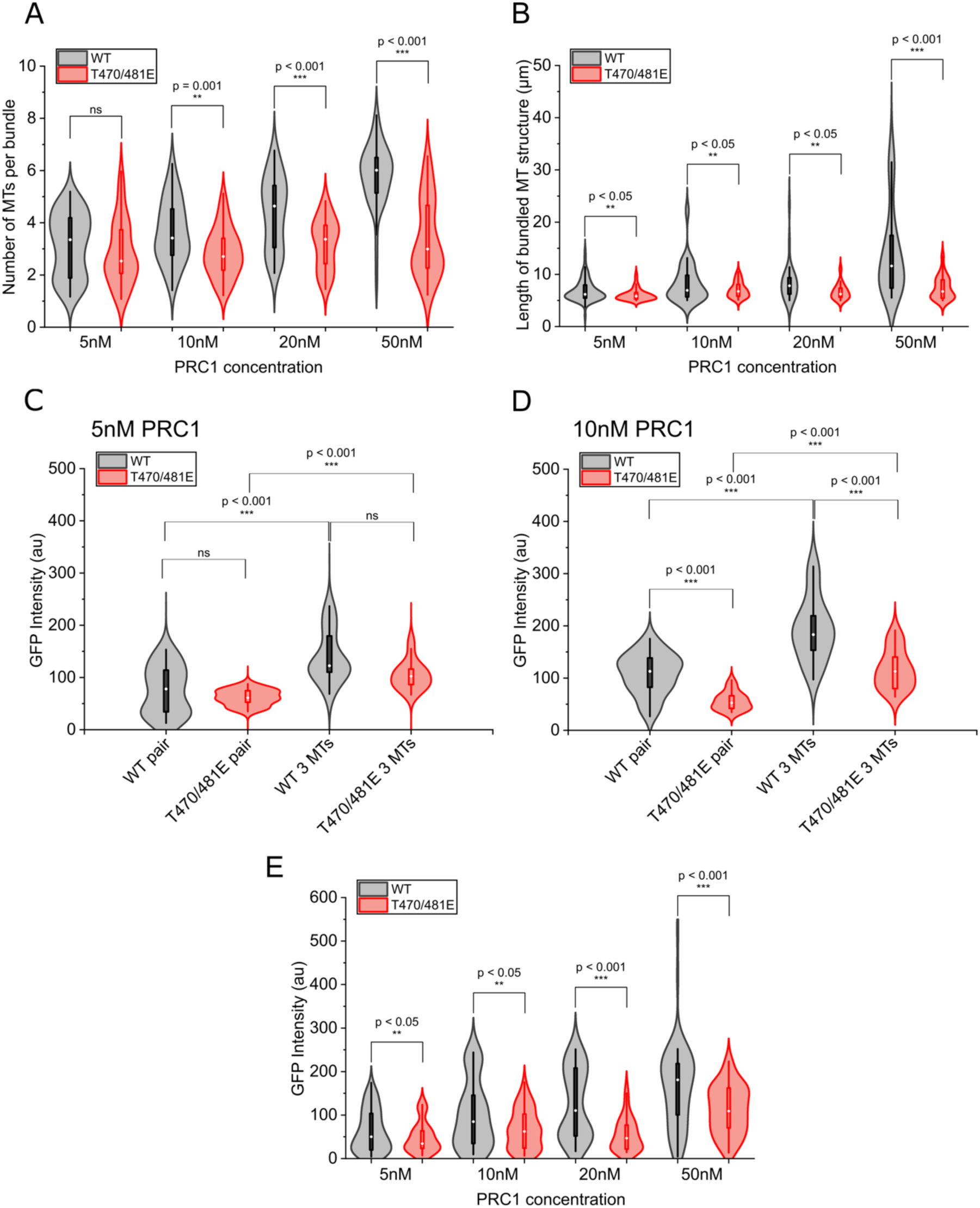
WT-PRC1 forms larger bundles containing more microtubules than T470/481E-PRC1. (A) Distributions of number of microtubules within individual bundles crosslinked by WT-PRC1 (black) and T470/481E-PRC1 (red) at 5, 10, 20, and 50nM PRC1 concentration (5nM: N = 63 WT, 52 T470/481E; 10nM: N = 54 WT = 60 T470/481E; 20nM: N = 38 WT, 42 T470/481E; 50nM: N = 50 WT, 50 T470/481E). (B) Distributions of lengths of microtubule bundle lengths when crosslinked by WT-PRC1 (black) and T470/481E-PRC1 (red) at 5, 10, 20, and 50nM PRC1 concentration. (C) Distribution of GFP-PRC1 intensity within pairs and bundles containing three microtubules for WT-PRC1 (black) and T470/481E-PRC1 (red) when formed with 5nM solution PRC1 concentration. (D) Distribution of GFP-PRC1 intensity within pairs and bundles containing three microtubules for WT-PRC1 (black) and T470/481E-PRC1 (red) when formed with 10nM solution PRC1 concentration. (E) Distribution of GFP-PRC1 intensity within microtubule bundles of any size for WT-PRC1 (black) and T470/481E-PRC1 (red) when formed with either 5, 10, 20, or 50nM solution PRC1 concentration.

We next sought to understand how PRC1 localization within microtubule pairs and bundles might differ between the two constructs. We first focused on bundles containing either two (“pairs”) or three (“triples”) microtubules. Linescan analysis of the GFP-PRC1 signal revealed that at low (5nM) concentrations of PRC1, statistically indistinguishable amounts of PRC1 are recruited to overlaps for both microtubule pairs and triples for both constructs (Fig. 2C). We next examined bundles assembled in the presence of 10nM PRC1, finding that the wild-type protein exhibited higher average signals within overlaps for both pairs and triples (Fig. 2D). Finally, we examined bundles of all sizes and observed a consistently higher recruitment of WT-PRC1 to overlaps than T470/481E-PRC1 across all solution concentrations of PRC1. Altogether, these data suggest that the formation of microtubule pairs is similar between the two PRC1 constructs, but that assembly of higher-order bundles containing three or more microtubules is enhanced for WT-PRC1 and diminished for T470/481E-PRC1.

### The formation of bundles occurs more rapidly with WT-PRC1 than T470/481E-PRC1

While the previous experiments revealed differences in the equilibrium distributions of crosslinked bundles formed, they did not provide information about the kinetic pathway by which bundling occurred. We therefore developed an assay which allowed us to monitor bundle formation in real time. Briefly, microtubules were introduced into a sample chamber with a PEGylated coverslip to prevent non-specific sticking. Methylcellulose (0.1% w/v) was introduced into the buffer which acted as a crowding agent and compressed the free microtubules into a quasi-2D plane near the coverslip, allowing for visualization by TIRF microscopy (Fig. 3A). Microtubules were free to diffuse in this plane and would often collide but rarely pass over one another. The open chamber allowed for direct addition of PRC1 molecules at 2.5 nM concentration.

**Figure 3.**
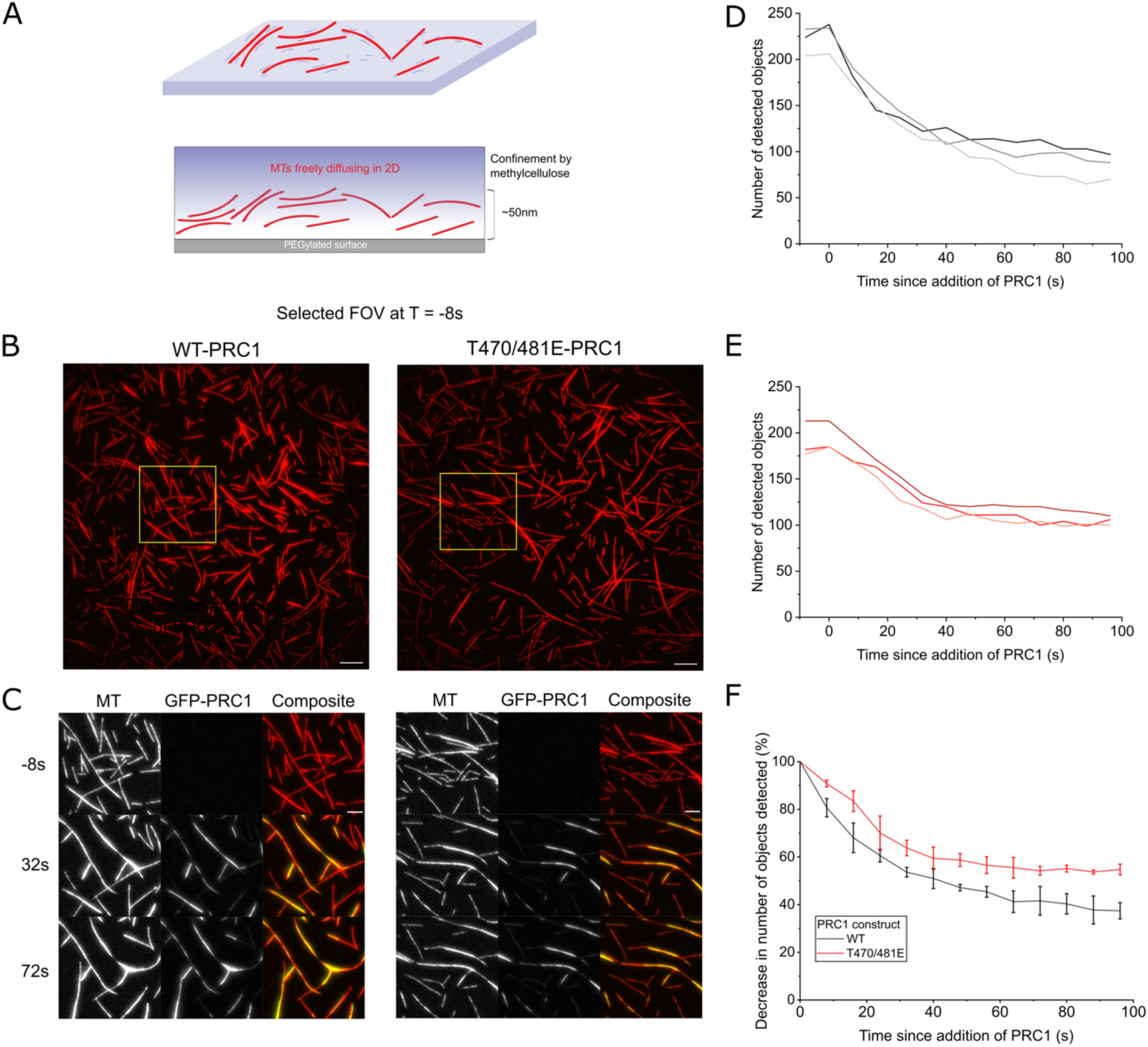
Formation of bundles occurs more rapidly with WT-PRC1 than T470/481E-PRC1. (A) Schematic depicting open-chamber assay with PEGylated surface, free microtubules, and added methylcellulose to confine free microtubules to the coverslip surface region for TIRF imaging. (B) Example field of view of microtubules taken 8 seconds before addition of PRC1. Scale bar = 10 µm. (C) Images of microtubules (red) and PRC1 (green) prior to (-8s), shortly after (32s), and long after (72s) addition of PRC1 to the sample chamber for both WT-PRC1 (left) and T470/481E-PRC1 (right). (D) The number of detected individual objects observed in the microtubule channel per field of view as a function of time for three different experiments using WT-PRC1. (E) The number of detected individual objects observed in the microtubule channel per field of view as a function of time for three different experiments using T470/481E-PRC1. (F) Average number of detected individual objects observed in the microtubule channel for both constructs as a function of time.

We imaged full fields (∼132x132 micron) and observed approximately 200 individual microtubules per field of view (Fig. 3B). Upon addition of GFP-PRC1 to the chambers, we observed a rapid increase in GFP signal within regions where two or more microtubules collided (Fig. 3C). Over time, as more microtubules diffused and came into contact, additional PRC1 would be recruited between these overlaps to form larger bundles. To quantify these data, we employed an object counting algorithm to detect individual structures. We performed this measurement for three different time lapse image series for both WT-PRC1 (Fig. 3D) and T470/481E-PRC1 (Fig. 3E). In both instances, we observed that as single microtubules began to crosslink into pairs and then further into higher order bundles, the number of detected objects decreased. In order to compare the two constructs directly, we normalized each trace by its initial object count, thereby reflecting a percentage decrease in total objects detected (Fig. 3F). Here, we observe a clear difference in the kinetics of higher-order microtubule bundling, with WT-PRC1 forming bundles more rapidly and plateauing at a much lower percentage than T470/481E-PRC1.

### Microtubule pairs bundled by T470/481E-PRC1 and WT-PRC1 both generate braking resistance against kinesin-driven sliding forces via a tip clustering mechanism

We recently reported on the behavior of crosslinked microtubule pairs disrupted by motor-protein induced forces, identifying two distinct modes of resistance to filament sliding^31^. To investigate how pair disruption under mechanical load might differ between T470/481E-PRC1 and WT-PRC1, we employed the same kinesin-driven microtubule gliding assay^31,40^. Briefly, a kinesin-1 (K439)^41^ construct was immobilized onto a passivated glass coverslip. Microtubules were incubated with 5 nM of PRC1 and were introduced into the flow chamber alongside either 100 µM or 2 mM ATP and in a buffer containing 0mM or 60mM KCl respectively. Upon contact with the kinesins, microtubule sliding began as the kinesins walked towards the plus-ends of microtubules, pushing the minus-ends outward and reducing the overlap length until the bundles fully separated. We observed a mixed population of single microtubules, pairs, and higher-order structures, and focused on pairs for further analyses.

After selecting microtubule pairs which were well isolated from other structures in the field of view and which exhibited overlaps of at least 2 microns in initial length, we generated kymographs to visualize the time course of bundle disruption (Fig. 4A, B). As the ends of the bundled microtubules are pushed in opposite directions via kinesin-1, the overlap length decreases. Upon disruption the slope changes, reflecting a change in velocity as the microtubules are no longer directly connected via PRC1 molecules. From the slope of these edges, we can calculate the change in velocity of microtubule pairs throughout disruption. We observed two velocity regimes per kymograph: one slower “bundled” velocity that occurred while microtubules were crosslinked, and a faster velocity after the microtubules “escaped” the bundle. We then calculated the ratio of bundled velocity (V_Bundled_) to escaped velocity (V_Escaped_). Using our previously reported criteria, we classified events with ratios greater than 0.4 as “braking” and those with ratios less than 0.4 as “coasting”^31^. We focused specifically on braking events and found no significant differences in the ratio distributions between the T470/481E and WT constructs under high ionic concentrations at saturating ATP conditions (Fig. 4C), nor at low ionic strength and low ATP concentration (Fig. 4E), suggesting that both PRC1 constructs could provide significant resistance against microtubule sliding.

**Figure 4.**
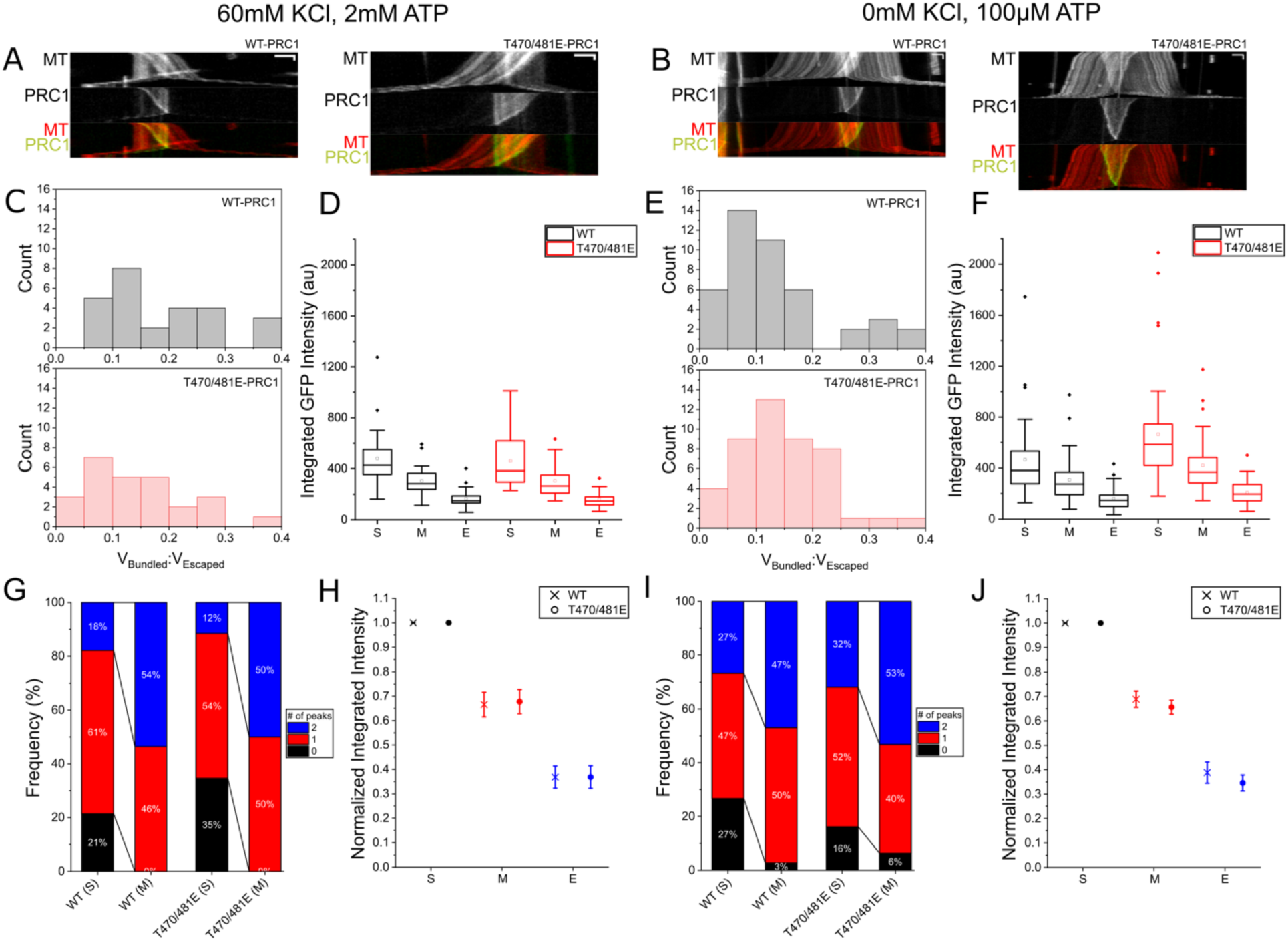
Both T470/481E-PRC1 and WT-PRC1 generate braking forces against kinesin-driven sliding forces via a tip clustering mechanism. (A) Representative kymographs of microtubule pairs crosslinked by WT (left) and T470/481E-PRC1 (right) being slid apart via kinesin-driven load in 60 mM additional KCl and 2 mM ATP (hereafter high salt/ATP. X-scale: 2 µm, Y-scale: 10 s). (B) Representative kymographs of microtubule pairs crosslinked by WT (left) and T470/481E-PRC1 (right) being slid apart via kinesin-driven load in 0mM additional KCl and 100 µM ATP (hereafter low salt/ATP. X-scale: 2 µm, Y-scale: 40 s). (C) Velocity ratio histogram of bundled to escaped velocities under high salt/ATP. WT: gray, N = 28; T470/481E: red. N = 26. (D) Box plots of overlap-bound GFP integrated intensity distribution within bundles at three overlap states (S = “start”, M = “middle”, and E = “end”) at high salt/ATP. Black = WT-PRC1, red = T470/481E-PRC1. Within each state, the GFP integrated intensity was statistically insignificant between the two constructs (p = 0.737, p = 0.974, and p = 0.756 for start, beginning, and end). (E) Velocity ratio histogram of bundled to escaped velocities in low salt/ATP. WT: gray, N = 45; T470/481E: red, N = 47. (F) Box plots of overlap-bound GFP integrated intensity distribution within bundles at three overlap states (S = “start”, M = “middle”, and E = “end”) at low salt/ATP. Black = WT-PRC1, red = T470/481E-PRC1. GFP integrated intensity was higher for T470/481E-PRC1 than WT-PRC1 at each state (p = 0.001, p = 0.007, and p = 0.014 for start, beginning, and end). (G) Distribution of detected peak number for WT (left) and T470/481E-PRC1 (right) at the start (S) and middle (M) of pair disruption at high salt/ATP. (H) Normalized GFP Integrated Intensity at high salt/ATP, values were not statistically significant between WT and T470/481E-PRC1 at the middle (p = 0.734) or end (p = 0.896) of disruption. (I) Distribution of detected peak number for WT (left) and T470/481E-PRC1 (right) at the start (S) and end (E) of pair disruption at low salt/ATP. (J) Normalized GFP Integrated Intensity at low salt/ATP, values were not statistically significant between WT and T470/481E-PRC1 at the middle (p = 0.142) or end (p = 0.122) of disruption.

Next, we focused on analyzing the distributions of PRC1 molecules via the GFP fluorescence signal throughout each disruption event. For each bundle, we considered three distinct points in the evolution of sliding: the initial distribution at full overlap length, an intermediate time point where the overlap length was reduced to half of its initial value, and the final state at short overlap just prior to bundle separation. We termed these overlap states “Start” (S), “Middle” (M), or “End” (E) and the integrated PRC1 intensity was measured for each bundle overlap for these three states (S4). At high ionic strength and saturating ATP, there was no significant difference in PRC1 recruitment and retention within overlaps between T470/481E and WT-PRC1 at each of the selected time points (Fig. 4D). However, at 100µM ATP and no additional salt in the system, a higher number of T470/481E-PRC1 molecules recruited within overlaps was observed for each timepoint (Fig. 4F). These findings suggest that a higher number of T470/481E-PRC1 molecules are required to form microtubule pairs to resist kinesin-driven sliding compared to WT-PRC1, further supporting our previous findings that T470/481E-PRC1 is less efficient at bundling compared to WT-PRC1.

We have previously shown that braking events correlate with the formation of dense PRC1 peaks near the microtubule ends. Therefore, we mapped the evolution of PRC1 peaks throughout bundle disruption by counting the number of detectable PRC1 peaks present within overlaps at early (S) and intermediate (M) time points. We found that these overlaps contained either an even distribution of PRC1 molecules with no distinct peaks, one peak at one end, or two peaks, one at each end. At saturating ATP and high salt, WT-PRC1 microtubule pairs most frequently had one peak present at the start, with fewer overlaps containing two peaks or no peaks (Fig. 4G). By the middle of bundle disruption, this population shifted towards a more even split of one peak or two peaks, but no overlaps contained zero peaks. Similarly, T470/481E-PRC1 microtubule pairs exhibited a distribution of mostly one peak present at the start of bundle disruption, with fewer overlaps containing two peaks or no peaks. Towards the middle of bundle disruption, this distribution shifted to an even split of one or two peaks, but no overlaps contained zero peaks. There was no statistically significant difference observed between the distribution of peak counts between WT and T470/481E-PRC1 at the beginning and near the middle of overlap disruption. We repeated the peak counting classification for PRC1-mediated microtubule pairs in the 100µM ATP and no additional salt condition. Here, we found similarly that there was no statistically significant difference in peak count distribution between the WT and T470/481E constructs at the start of disruption (Fig. 4I). Likewise, near the end of pair disruption, the distribution of peaks within microtubule pairs was similar for WT and T470/481E-PRC1. Together these results suggest that PRC1 clustering near tips occurs via similar pathways for both constructs, and leads to similar braking behavior in sliding bundles.

Finally, we calculated the normalized integrated intensity of PRC1 within the bundle overlaps examined in Figures 4D and 4F to determine how the percentage of PRC1 molecules lost during sliding might vary between the two proteins. The normalized integrated intensity was comparable between microtubule pairs bundled by WT and T470/481E-PRC1 both in high ionic conditions under saturating ATP (Figure 4H) as well in the absence of additional salt and under low ATP conditions (Figure 4J). Taken all together, these results suggest that for microtubule pairs crosslinked by PRC1, both the T470/481E and WT constructs can cause a significant slowdown of kinesin-driven sliding. For both proteins, this braking mechanism is linked to dense PRC1 peaks which form in response to external sliding forces near microtubule tips. Though it takes more T470/481E molecules within the overlap to form pairs that enter into a braking mode, the change in distribution and loss of molecules follows a similar pathway to the WT-PRC1 construct.

### Higher-order microtubule bundles are disrupted faster and more completely when crosslinked by T470/481E-PRC1 than WT-PRC1

We next sought to determine the pathways by which higher-order microtubule bundles could be disrupted by kinesin-mediated forces. We generated crosslinked bundles in the presence of 20nM PRC1 and introduced these bundles to our kinesin disruption assay. We then selected larger bundles which contained at least six microtubules for further analysis. We observed that over the course of approximately three minutes, WT-PRC1 bundles were disassembled slowly with frequent loss of single and paired microtubules from the larger structure (Fig. 5A). Over the same time course, T470/481E-PRC1 bundles were pulled apart much faster and frequently were fully disassembled after only a few minutes, with the most common rupture pathway being stochastic loss of single microtubules (Fig. 5B). To quantify these behaviors, we selected N = 15 bundles containing at least 6 microtubules each per PRC1 construct and protein concentration and determined the extent to which they reduced in size as they were disassembled. Bundles assembled in the presence of 20nM WT-PRC1 disrupted significantly more slowly than those assembled with T470/481E-PRC1 (Fig. 5C). When we performed the experiments using 50nM solution PRC1 concentration, we observed that WT-PRC1 bundles almost never disrupted, remaining intact even under significant sliding forces generated by kinesin. In contrast, T470/481E-PRC1 mediated bundles exhibited substantial disruption, often losing single or paired microtubules in a matter of minutes. Together, these data reveal that WT-PRC1 higher order bundles are much more stable and resistant to forces that would slide the microtubules apart, whereas T470/481E-PRC1 bundles can readily be slid apart under motor protein driven forces.

**Figure 5.**
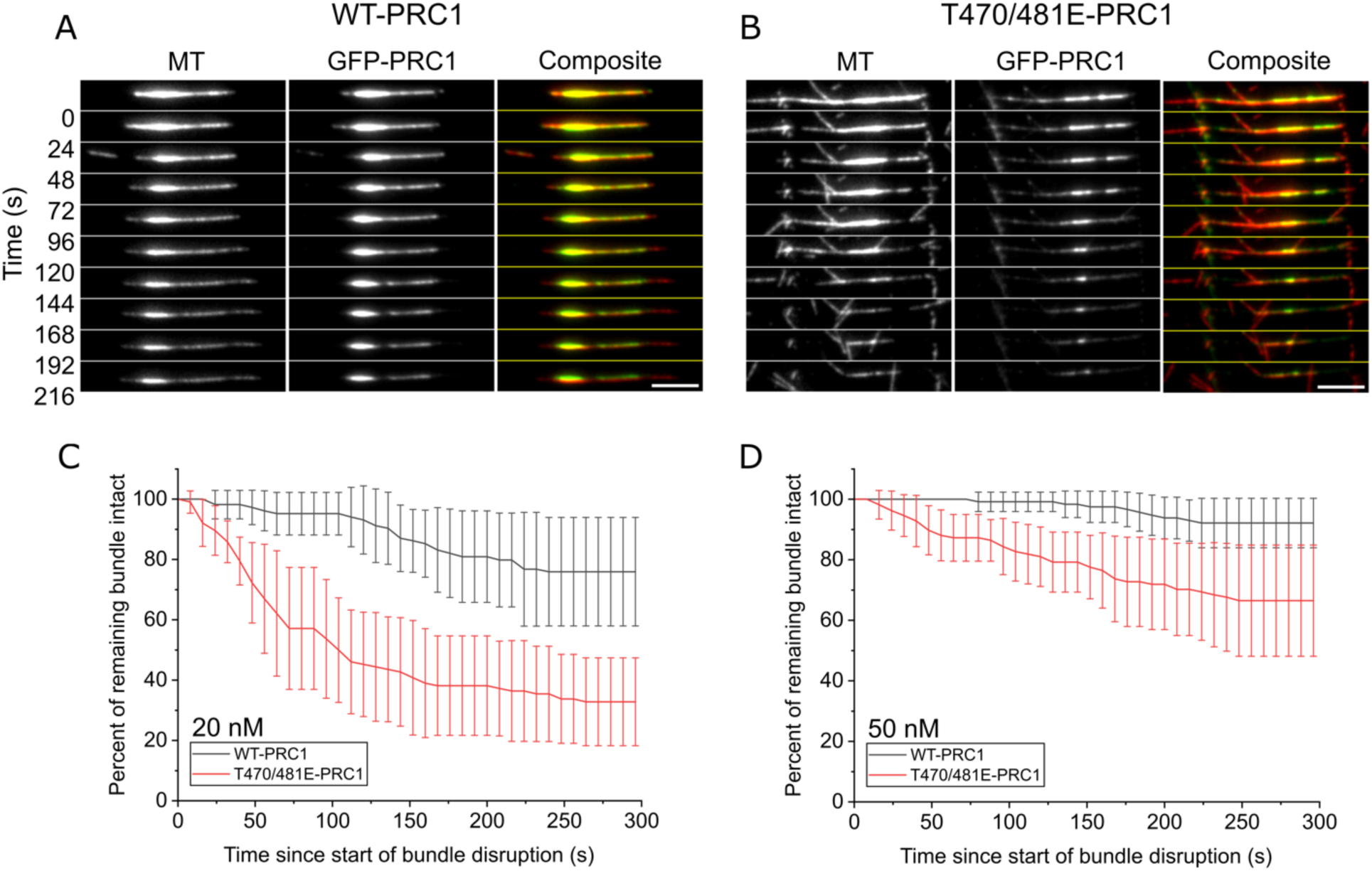
Higher-order microtubule structures crosslinked by T470/481E-PRC1 are disrupted more rapidly and completely compared to WT-PRC1. (A-B) Time-lapse images of higher-order structures crosslinked by WT-PRC1 (left) or T470/481E-PRC1 (right) imaged by TIRF. Left: microtubules are shown. Middle: PRC1 is shown. Right: composite is shown (microtubules in red, WT-PRC1 in green). Time elapsed in seconds is shown on the left axis. Scale bar = 5 µm. (C) Time course of kinesin-driven disruption in higher-order (6-8 microtubule) structures bound by 20 nM PRC1. Structures assembled by T470/481E-PRC1 (red) are disassembled significantly more quickly compared to structures assembled by WT-PRC1, with fewer microtubules remaining in the final observed structure (t-test information shown in S6). (D) Time course of kinesin-driven disruption in higher-order (6-8 microtubule) structures bound by 50 nM PRC1. Structures assembled by T470/481E-PRC1 (red) are disassembled significantly more quickly compared to structures assembled by WT-PRC1 (t-test information shown in S6). Bundles crosslinked by WT-PRC1 at 50 nM are rarely perturbed by kinesins, but T470/481E-PRC1 bundles are slightly disrupted throughout imaging.

## Discussion

We have investigated the differences between dephosphorylated WT-PRC1 (which reflects its anaphase biochemical state) and a phosphomimetic T470/481E-PRC1 (reflecting a metaphase state) in the context of building microtubule structures both at equilibrium as well as in response to disruptions under mechanical load. We find that WT-PRC1 and T470/481E-PRC1 behave similarly when building and maintaining microtubule pairs against motor-driven microtubule sliding. In contrast, bundles formed using T470/481E-PRC1 are shorter and contain fewer microtubules, form at a slower rate, and are more susceptible to disruption by sliding forces than WT-PRC1.

How could two amino acid residue substitutions that mimic phosphorylation at sites adjacent to, but not directly within, the microtubule binding domain propagate to influence micron-scale properties of bundle assembly? To provide a framework for addressing this question, we performed protein structure predictions using AlphaFold2 for either full-length WT or T470/481E-PRC1 bound to an αβ tubulin dimer^39^. The predicted structure and position relative to the αβ tubulin of PRC1’s microtubule-binding spectrin domain (Fig. 6A, purple) were in very good agreement with that of a published structure acquired via cryo-EM (Fig. S4)^24^. In contrast, small changes in the positioning of the spectrin domain, including a slight tilt and rotation relative to the tubulin surface, were predicted for T470/481E-PRC1 (Fig. 6A, yellow). We also noted that the interaction sites between the αβ tubulin dimer and residues 470-481 within PRC1 were shifted due to the two amino acid substitutions at sites 470 and 481 (Fig. 6B). Together, these predictions suggest that introduction of negative charge within regions of the protein adjacent to the microtubule binding domain can induce local conformational changes within PRC1 and between its interaction sites at the microtubule surface. Interestingly, our data suggests that these subtle differences could perhaps propagate to influence crosslinking behaviors across the full length of the PRC1 dimer and perhaps even within ensembles of multiple PRC1 molecules.

**Figure 6.**
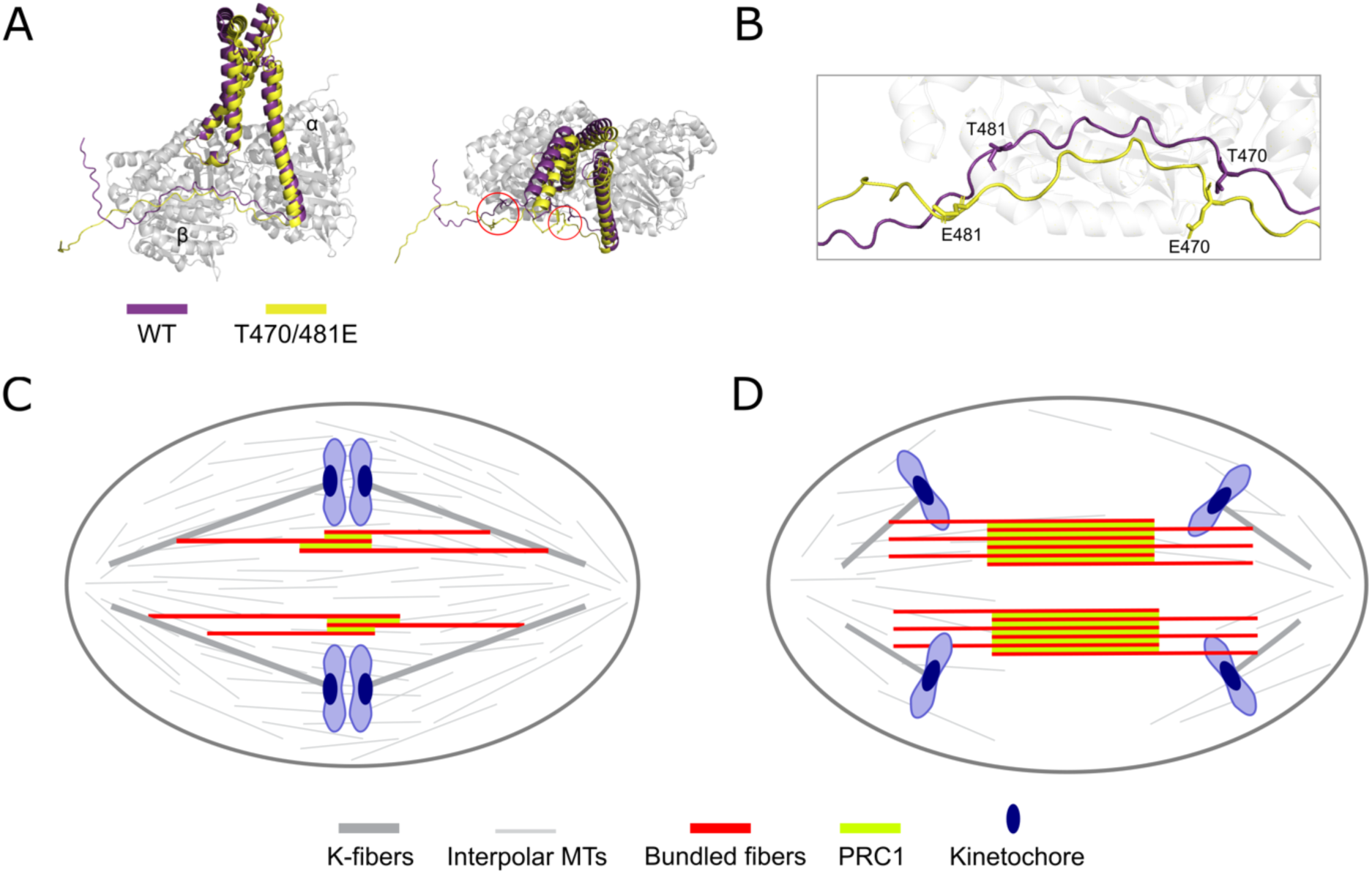
Structural predictions and proposed model for bundle arrangement via PRC1 in metaphase and anaphase. (A) AlphaFold2 prediction of full-length PRC1 bound at the α-β tubulin dimer interface (white). The spectrin domain and first residues of the coiled-coil region of T470/481E-PRC1 (yellow) are shown alongside WT-PRC1 (purple). Residues 320-500 are shown for each PRC1 construct prediction. The alpha and beta tubulin chains are indicated as “α” and “β”, respectively, and shown in gray. Alignment of the two predicted structures at their α-β chains yielded RMSD = 0.299. (B) Inset of AlphaFold2 prediction showing residues in WT-PRC1 that were altered using phosphomimetic substitution to create T470/481E-PRC1. (C-D) Proposed schematic of mitotic spindle bundles organized by PRC1 in metaphase (C) and anaphase (D). Kinetochore fibers are shown in gray, unbundled interpolar microtubules in light gray, bundled interpolar microtubules in red, PRC1 in green, and chromosomes in blue.

What could the consequences of differential microtubule bundling be in dividing cells? We hypothesize that if PRC1 is phosphorylated by CDK/cyclin-B at threonine residues 470 and 481 during metaphase, it would organize smaller microtubule structures that are more compliant against internal sliding forces and pushing/pulling forces when engaged in bridging of sister kinetochore fibers (Fig. 6C). This mechanical organization would likely be amenable to providing both structural stability and flexibility when subject to the bidirectional oscillations during chromosome alignment at the metaphase plate^9,11,12^. Upon dephosphorylation of PRC1 at anaphase onset, bundles would be more likely to grow in diameter with the stable addition of microtubules, and they would provide enhanced resistance against poleward forces^30,31,42^ (Fig. 6D). A larger number of microtubules stabilized by a highly dense packing of PRC1 molecules later in anaphase would provide a strong mechanical resistance to force and favor the organization of a stable spindle midzone^22,38,42^.

Regulation of microtubule interactions by phosphorylation has been well studied in the context of kinetochore-microtubule attachments. For example, phosphorylation of the Ndc80 complex by the Aurora B kinase loosens microtubule interactions and is linked to the regulation of chromosome motions and error correction processes^43,44^. In contrast, our understanding of how proteins that organize interpolar microtubule networks are differentially regulated by phosphorylation and dephosphorylation events is more limited. In addition to PRC1, crosslinking motor proteins like kinesin-5 are known to be targeted by kinases, particularly within their C-terminal domains^45–47^. How these PTMs modulate key parameters such as production of sliding forces, velocities, and cooperativity between motors within overlaps will need to be carefully examined. It is likely that understanding the coordination between motor and non-motor MAPs at distinct phases of mitosis will give insights into the biophysical rules governing spindle assembly.

## Acknowledgments

We would like to thank Dr. Susan Gilbert (RPI) for the gift of the kinesin-1 K439 plasmid and Dr. Blanca Barquera (RPI) for assistance with construct design. We also wish to thank Kristen Hewes, Jacob Palumbo, Drs. Susan Gilbert and Marvin Bentley (RPI) for providing advice and suggestions for experiments and data analysis methods. We also wish to thank members of the Forth lab for helpful discussions and critical reading of the manuscript. This work was supported by startup funds provided by the School of Science at Rensselaer Polytechnic Institute, NIH NIGMS R01GM149782 and NSF 2153374 to S.F., and T32AG057464 and T32AG078123 to E.T.

## Author contributions

E.T., A.A., and S.F. conceived of experimental design. E.T. and A.H. performed data acquisition and analysis. E.T. and A.A. performed expression and purification of proteins used in the study. E.T. and G.S. performed structural prediction analysis. E.T. and S.F. wrote and edited the manuscript. S.F. secured funding to perform the research.

