## Supplemental Information for "Insights into the role of phosphorylation on microtubule crosslinking by PRC1"

### Supplemental Figures and Information to accompany “Insights into the role of phosphorylation on microtubule crosslinking by PRC1”

Ellinor Tai, Austin Henglein, Angus Alfieri, Gauri Saxena, Scott Forth*

**Supplemental Figures S1-S4**

**Figure S1: Linescans of rhodamine fluorescence signal within individual objects is used to calculate the number of microtubules within structures**

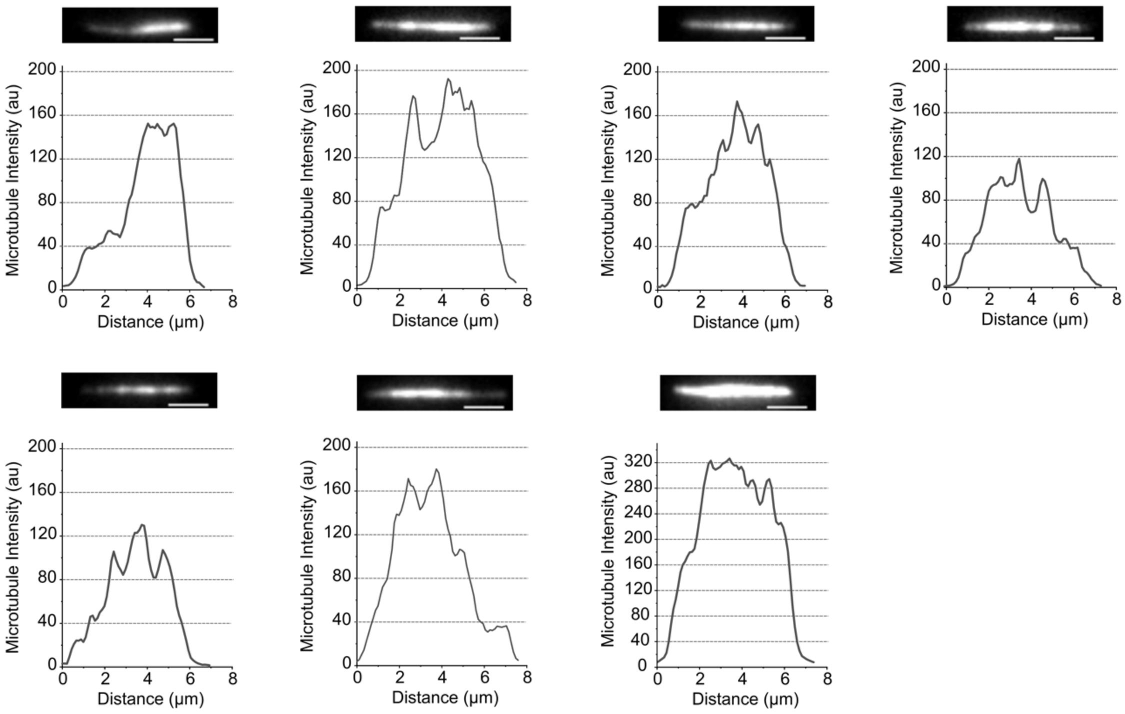


**Figure S1** Example images of microtubule bundles (rhodamine channel, grayscale) and the corresponding line scan profiles along their lengths. Dashed lines represent the intensity signal from a single microtubule (~40 a.u.). Scale bars = 2 μm.

**Figure S2: Example of PRC1 peak detection within microtubule pairs being disrupted under kinesin-driven load**

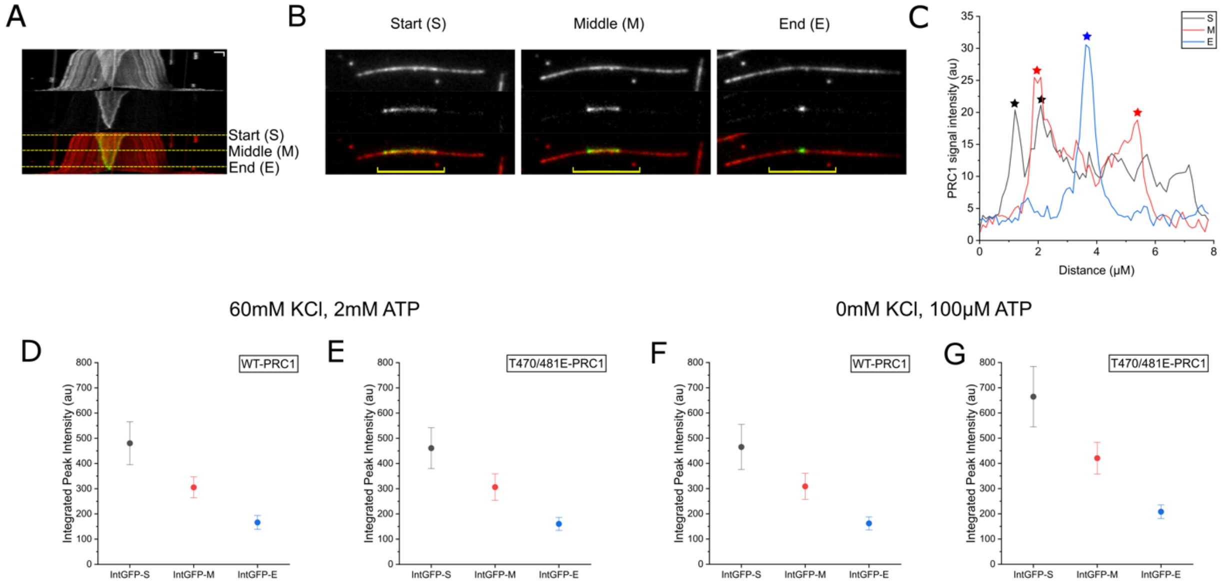


**Figure S2** (A) Representative kymograph of a microtubule pair being ruptured under the activity of surface-bound kinesin motor proteins (top: microtubule, middle: PRC1, bottom, composite). Dashed lines indicate time points used for analysis: Start (S) reflects the initial observed long overlap length, Middle (M) reflects a reduction in overlap length by half, and End (E) reflects a point just prior to complete disruption. X-scale: 2 μm, Y-scale: 40 s. (B) Images of a disrupting bundle at time points S, M, and E (top: microtubule, middle: PRC1, bottom, composite). The yellow line represents the length of the region selected for intensity profile analysis (8 μm). (C) Intensity profiles from (B) for time points S, M, and E. Stars indicate detected peaks. (D-G) Average absolute integrated intensity values for (D) WT-PRC1 and (E) T470/481E-PRC1 at high salt/ATP and (F) WT-PRC1 and (G) T470/481E-PRC1 at low salt/ATP.

**Figure S3: Tracking disruption of individual higher-order structures over time reveals that T470/481E-PRC1 bundles disrupt more completely and rapidly**

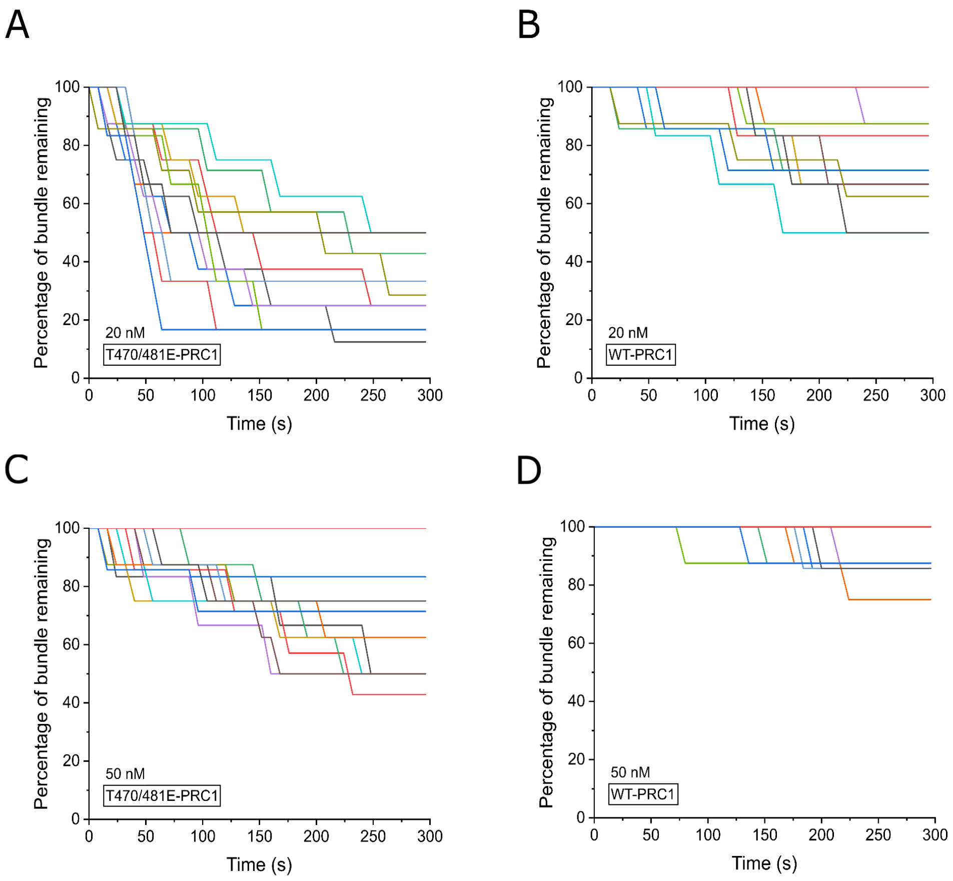


**Figure S3** (A-D) Individual large bundle disruption event trajectories for bundles crosslinked by PRC1, plotted as a percentage of microtubules remaining relative to the initial observed structure for (A) 20 nM T470/481E-PRC1, (B) 20 nM WT-PRC1, (C) 50 nM T470/481E-PRC1, and (D) 50 nM WT-PRC1. N = 15 bundles per each construct and condition.

**Figure S4: AlphaFold2 prediction of the spectrin domain of full-length WT-PRC1 aligned to published cryo-EM structure PDB: 5KMG**

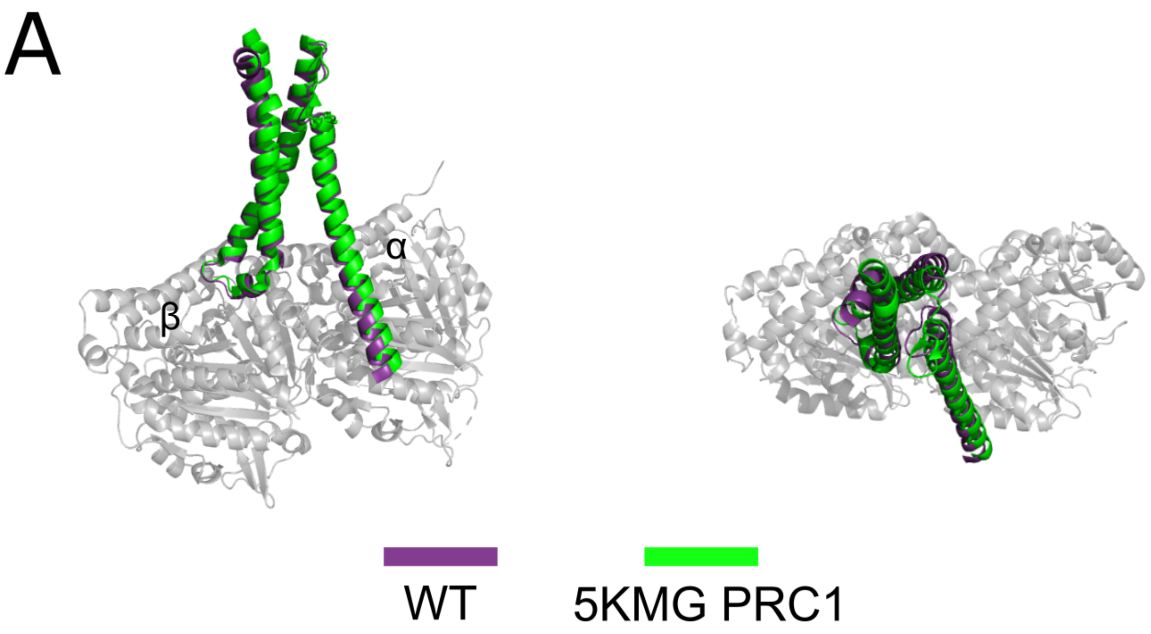


**Figure S4** Comparison of AlphaFold2 prediction of PRC1's microtubule binding domain (purple) and published cryo-EM structure 5KMG (green). The alpha and beta tubulin chains are indicated as "α" and "β", respectively, and shown in gray. The predicted structure was aligned to the published structure for verification of similarity (RMSD = 1.317Å).

#### Materials and Methods

##### Protein Expression and Purification

*WT-PRC1*

Full-length human PRC1 Isoform 2 was cloned into a pET-DUET plasmid containing an N-terminal histidine tag followed by a Tobacco Etch Virus (TEV) cleavage site. An eGFP sequence was inserted between the TEV cleavage site and the N-terminus of PRC1 with a 'AAA' linker sequence just after the eGFP. Proteins were expressed via BL21(DE3) Rosetta *Escherichia coli* cells (Novagen). Full-length PRC1 was expressed for 3.5 hours at 18$^{o}$C after induction with 0.5 mM IPTG. Cells were lysed via sonication in lysis buffer (1 mg/mL lysozyme, 50 mM phosphate (pH 8.0), 10 mM imidazole, 1% Igepal and HALT protease inhibitor (Pierce), 2 mM TCEP Bond Breaker, 2 mM Benz-HCl, 1 mM PMSF). The lysate was clarified by ultracentrifugation and the supernatant was incubated with Ni-NTA for 1 hour at 4$^{o}$C (G Biosciences). The resin was then washed with Wash Buffer (50 mM phosphate, (pH 8.0), 500 mM KCl, 10 mM imidazole, 0.1% tween, 0.5 mM TCEP, 1 mM PMSF) and the protein was then eluted using Elution Buffer (50 mM phosphate (pH 7.0), 250 mM imidazole, 150 mM KCl, 0.5 mM TCEP). Elute was pooled and concentrated to a volume of approximately 1 mL. Next, 1/30 w/w Pro TEV protease (PROMEGA), 1 mM DTT, and 50 uL of Pro TEV 20X buffer were added and the sample incubated in a 30˚C water bath for 15 minutes before overnight dialysis at 4$^{o}$C in Gel Filtration Buffer (1X BRB80 (pH 6.8), 150 mM KCl, 10 mM bME). Size exclusion chromatography (Superose-6 increase 10/300 column, GE Healthcare) was then performed in Gel Filtration Buffer with a Shimadzu High Performance Liquid Chromatograph. After collecting peak fractions and concentrating to ~0.5 mg/mL, sucrose was added to 35% w/v before flash freezing in liquid nitrogen.

*T470/481E-PRC1*

Sites Thr-470 and Thr-481 in full-length PRC1 Isoform 2 were mutated to glutamates using the QuikChange Site-Directed Mutagenesis Kit (Agilent) to generate the phosphomimetic construct, which was expressed and purified with the same protocol used for WT-PRC1.

*Kinesin-1 K439*

Plasmid encoding a truncated kinesin-1 construct (K439) with a C-terminally fused EB1 sequence to create stable dimers and a $His_8$ tag[41] was generously donated by the Dr. Susan Gilbert Lab. The protein expression protocol was identical to PRC1, except induction occurred overnight for 18 hours at 16$^oC$. The pellet was resuspended in lysis buffer (10 mM $NaPO_4$ (pH 7.2), 300 mM NaCl, 2 mM $MgCl_2$, 0.1 mM EGTA, 10 mM PMSF, 1 mM DTT, 0.2 mM ATP, 1 mg/mL lysozyme, 30 mM imidazole, 3 uL benzonase (Novagen)) and sonicated. The lysate was clarified by ultracentrifugation and the supernatant incubated with Ni-NTA for 1 hour at 4$^oC$ (G Biosciences). The resin was then washed with Wash Buffer (20 mM $NaPO_4$ (pH 7.2), 300 mM NaCl, 2 mM $MgCl_2$, 0.1 mM EGTA, 0.02 mM ATP, 50 mM Imidazole) and the protein was then eluted using Elution Buffer (20 mM $NaPO_4$ ( pH 7.2), 300 mM NaCl, 2 mM $MgCL_2$, 0.1 mM EGTA, 1 mM DTT, 0.02 mM ATP, 400 mM imidazole). Elute was pooled and concentrated to a volume of approximately 1 mL. The protein was then dialyzed overnight at 4$^oC$ in Dialysis Buffer (20 mM HEPES (pH 7.2), 300 mM NaCl, 0.1 mM EGTA, 0.1 mM EDTA, 5 mM MgAc, 50 mM KAc, 1 mM DTT). The next day the protein was dialyzed for 1 hour at 4$^oC$ in Buffer 1 (20 mM HEPES (pH 7.2), 200 mM NaCl, 0.1 mM EGTA, 0.1 mM EDTA, 5 mM MgAc, 50 mM KAc,1 mM DTT) followed by dialysis for 1 hour at 4$^oC$ in Buffer 2 (20 mM HEPES (pH 7.2), 150 mM NaCl, 0.1 mM EGTA, 0.1 mM EDTA, 5 mM MgAc, 50 mM KAc,1 mM DTT) Further purification was performed using size exclusion chromatography (Superose-6 increase, GE Healthcare) using Shimadzu High Performance Liquid Chromatograph. Peak fractions were pooled and dialyzed for 3 hours at 4$^oC$ in Buffer 3 (20 mM HEPES (pH 7.2), 100 mM NaCl, 0.1 mM EGTA, 0.1 mM EDTA, 5 mM MgAc, 50 mM KAc,1 mM DTT, 5% Sucrose). Protein was flash frozen in liquid nitrogen.

**Microtubule Preparation**

Microtubule tubulin reagents were purchased from Cytoskeleton, Inc. Microtubules for surface-immobilization were generated via mixture of rhodamine tubulin (TL590M) and unmodified tubulin (T240) at a ratio of 1:20 along with 1 mM GMPCPP. Microtubules were polymerized at 37$^{o}$C for 75 minutes before clarification and stabilization in 20 µM Taxol following published protocols[48].

**Static Bundling and Kinesin-driven Gliding Assay Flow Chamber Construction:**

The flow chamber design and assay preparation were modified from a previously described protocol[31]. Anti-parallel microtubule bundles were constructed using passivated glass coverslips coated with SVA-PEG at a ratio of 50 PEG:1 biotin-PEG. All reagents were prepared with 1X BRB80 buffer. PRC1 (5-50 nM) was mixed with microtubules and allowed to incubate while the chamber was being constructed. Following each reagent flow-in and incubation, a flush with ~3 chamber volumes of 1X BRB80 was performed. Reagents were introduced stepwise with the following order and incubation times: (1) 0.5 mg/mL neutravidin for 2 minutes; (2) 0.5 mg/mL alpha casein 5 minutes; (3) 0.01 mg/mL Anti-Histidine Antibody for 5 minutes; (4) 0.25 µg/mL K439 for 5 minutes; (6) PRC1-microtubule bundles were added to the final Reaction Buffer (0.5 mg/mL alpha casein, 20 µM Taxol, 1.25 mM $MgCl_2$, 0.125 mM EGTA, Oxygen Scavenging System (4.5 mg/mL glucose, 350 U/mL glucose oxidase, 34 U/mL catalase, 1mM DTT), 100-2000 µM ATP. The chamber was then sealed with clear nail polish prior to experiments.

**Real-time Bundle Formation Chamber Construction:**

Microscope cover glass slips (22 mm x 60 mm) were treated with the following reagents and times, with a dd$H_2O$ rinse between each step: 100% methanol for 5 minutes; HPLC-grade acetone for 5 minutes; 2% Micro90 solution for 20 minutes; 0.1M NaOH solution for 20 minutes, then stored in 100% ethanol overnight. Slides were then treated with 0.01

mg/mL Vectabond solution for 5 minutes prior to chamber assembly. A 1.0mm diameter silicone o-ring was placed onto the center of each 22 mm x 60 mm cover glass slip and gently pressed on to create a seal between the glass and o-ring. A layer of SVA-PEG was pipetted into each o-ring and sealed with a glass coverslip. Slides were then placed in a humidity chamber on a gel rocker set to gentle speed for 40 minutes. The slides were then removed from the humidity chamber, the coverslips sealing the o-rings removed. SVA-PEG was pipetted out of each o-ring, and replaced 3 times with 1X BRB80 to wash. Each chamber was then dried by gently flipping the slides o-ring-side down onto a KimWipe, then used within 6-8 hours after drying.

Reagents were introduced stepwise with the following order and incubation times: (1) 0.5 mg/mL alpha casein for 5 minutes followed by rinsing with 1X BRB80; (2) microtubules in Crowding Assay Buffer with final volume 40 uL and w/v solution of 0.1% methylcellulose (0.5 mg/mL alpha casein, 20 μM Taxol, 1.25 mM $MgCl_2$, 0.125 mM EGTA, Oxygen Scavenging System (0.5 mg/mL alpha casein, 20 μM Taxol, 1.25 mM $MgCl_2$, 0.125 mM EGTA, Oxygen Scavenging System (4.5 mg/mL glucose, 350 U/mL glucose oxidase, 34 U/mL catalase, 1mM DTT) for 5 minutes; (3) additional 40 uL 0.1% methylcellulose Crowding Assay Buffer, then immediately placed onto the scope for imaging.

Using a extended tip 10 uL pipette tip, 2-4uL of 100 nM WT or T470/481E-PRC1 (final total chamber concentration 2.5-5 nM) was added into the o-ring containing the microtubule mix within the first 30 seconds of image acquisition by gently depressing the plunger button before the tip of the pipette touched the microtubule mix so that a droplet of PRC1 emerged, allowing for capillary action to merge the PRC1 droplet and microtubule mix contained in the o-ring.

**Structure Prediction:**

Google Colab AlphaFold2 was used to predict the structure of monomeric full-length human PRC1 Isoform 2 bound at its microtubule binding domain to an αβ tubulin dimer[39]. The FASTA file sequence chains for the published cryo-EM structure of PRC1-SC bound to the microtubule[24] (RCSB PDB ID: 5KMG) were used as the template for the query sequence of full-length WT-PRC1 bound to the microtubule. We replaced the PRC1-S sequence residues (containing the full spectrin domain, residues 345-467) with the sequence for full-length monomeric PRC1[49] (RCSB PDB ID: 4L3I, residues 1-620). To generate the query sequence for full-length monomeric T470/481E-PRC1 bound to the microtubule, we repeated the same process as above and changed the threonines at 470 and 481 to glutamic acid. For both query sequences, the αβ tubulin chains were not altered. Protein structure predictions were run on ColabFold v1.5.5 (AlphaFold2 using MMseqs2). PDB100 was used as the database for templates. Runs were performed using alphafold_multimer_v3 for complex prediction, and a separate MSA was run for each protein chain. The default Colab AlphaFold2 settings were used for other parameters. The number of recycle for each complex predicted was 3. Visualization of the predicted structure was done using PyMOL. Agreement between our predicted full-length WT-PRC1-MT structure and the published structure was assessed by using the alignment plugin in PyMOL to align our prediction with the PDB file for PRC1-S (5KMG), which also calculated an RMSD value. The αβ tubulin chains of both full-length WT-PRC1 and T470/481E-PRC1 were aligned to show the positioning of the spectrin domain for each construct. Only residues 320-500 in both PRC1 chains were shown in the models.

#### IMAGE QUANTIFICATION AND STATISTICAL ANALYSIS

##### Image Acquisition:

Microtubule bundles were imaged using two-channel TIRF microscopy using the following laser lines and exposure times: GFP-tagged PRC1 molecules were excited using a 488 nm laser (60% power, 100 ms exposure); and rhodamine microtubules were excited using a 561 nm laser (30% power, 100 ms exposure). Images were acquired using a Photometric Prime 95B camera controlled with Nikon NIS Elements software at overall acquisition rates of one frame per ~5 seconds or one frame per ~2 seconds. For microtubule pair analysis, images were visually screened to ensure that there were only two microtubules per bundle and there were no additional interactions with microtubules from other bundles.

##### Image Analysis

Analysis of fluorescent data and generation of intensity linescan data sets were performed using FIJI (ImageJ)[50] tools and Microsoft Excel. In all analyses performed, a linescan width of 5 pixels was used. In the static equilibrium experiments, a linescan was drawn along the length of each bundle in FIJI and the fluorescence intensity data along this line recorded for both the microtubule and PRC1 channels.

For pair rupture experiments, a linescan was drawn along the entire path of bundle separation in FIJI and the fluorescence intensity data along this line was saved for each frame, resulting in a 2D array of intensity values at all recorded time points. Kymographs were generated using the KymoResliceWide[51] plugin in FIJI. The “Start” (S) rupture timepoint for each pair was identified as the time at which overlaps were longest in length prior to disruption. “Middle” (M) timepoints were identified by tracking pair disruption until the overlap was 0.5 times its maximum length at S. “End” (E) timepoints were identified as the frame prior to complete separation of the bundle, where overlaps were the shortest. A peak detection algorithm was employed via a custom Excel script and used to find changes in slope in the GFP channel at these classified timepoints. An empirically determined maximum peak width value was used to filter the results and eliminate false peaks. After timepoint classification and peak identification, a summation over GFP intensity values in this region to determine integrated signal intensity could then be performed.

To identify the number of objects per given timepoint in live bundle formation assays, the 3D Objects Counter[52] plugin in FIJI was used. Substacks were taken of the rhodamine microtubule channel for each individual timepoint analyzed within a video to generate individual images of the frames and smoothed to increase object detection accuracy. The size filter in 3D Object Counter was set to the default maximum and a minimum of 100 to filter out smaller artifacts. The threshold minimum for each frame analyzed was set so the preview outlined microtubule structures as tightly as possible without causing any gaps. The resulting object count value per frame was then recorded for plotting.

**Statistical Analysis**

Datasets were compared using the Student's t-test function in OriginPro or Fisher's exact test in GraphPad. All relevant figure legends and sample sizes are described in the figure captions.
